# A New Computational Inverse Modeling Approach for Cellular Traction Force Microscopy that Accounts for Hydrogel Compressibility

**DOI:** 10.1101/2025.07.03.663064

**Authors:** Gabriel Peery, Toni M. West, Sanjana S. Chemuturi, Jodie H. Pham, Giovanni Ferrari, Michael S. Sacks

## Abstract

We have recently documented significant compressible behaviors in hydrogels implemented in 3D traction force microscopy (TFM). Therefore, here we have developed a new computational pipeline that accounts for this observation. Additionally, the new method accurately recovers large ranges and spatial heterogeneity of hydrogel moduli induced by cellular remodeling associated with enhanced extracellular matrix secretion and MMP-degradation. The algorithm sought best fit of the 3D displacement field with a multi-stage approach, wherein the Tikhonov regularization parameter in L-BFGS was progressively lowered. Forward simulations were performed in FEniCS, with gradients computed with FEniCS-adjoint and MOOLA to weight degrees of freedom according to hydrogel volume affected. Once developed, we conducted a series of synthetic test cases applying actual cell geometries, experimentally-matched compressibility, and realistic displacements with experimental noise levels. Employing an incompressible material model resulted in predicted moduli with over 415% mean relative error and predicted strain energies 5-fold greater than the prescribed values. Moreover, errors in predicted traction forces were amplified by a factor of 10. Thus, accounting for hydrogel compressibility was critical for accurate hydrogel moduli and strain energy recovery. To demonstrate the utility of our approach, we applied it to TFM data of human mitral valve interstitial cells embedded in PEG hydrogels with pre-altered moduli of 54 Pa. We determined that *J* ∈ [0.45, 1.66] and local hydrogel moduli exhibited large variations, 3.6 Pa to 2.4 MPa. This study underscores the need for correct handling of hydrogel compressibility for accurate estimation of local hydrogel moduli and traction forces.

## 1. INTRODUCTION

Traction force microscopy (TFM) is a powerful, well-established experimental-computational approach for estimating cellular contractile forces. In 3D TFM, cells are integrated into hydrogels that emulate certain aspects of the cell’s natural micro-environment and contain embedded micro-spheres that act as fiducial markers to track local displacements, typically before and after chemical treatment to modify cellular force generation. The resulting kinematic data by itself can be helpful to assess cellular contractile behaviors and cellular shape changes [1–4]. Additional mechanical quantities, such as cellular surface traction forces and hydrogel strain energy density fields, are also determined as they are better reflections of cellular biophysical activity [3].

Regardless of the details of the approach, TFM analyses rely on a detailed knowledge of the local cellular displacement field and hydrogel mechanical behaviors. Popular approximations for modeling hydrogel mechanical behaviors include treating the system as 2D [5, 6] or using a linear elastic material model [5–8], but these simplifications are not appropriate for capturing cellular behavior on 3D substrates with large deformations. Without these simplifications, large deformation TFM inverse analyses must address challenges involved in achieving the necessary numerical stability, accounting for ill-posedness of the nonlinear inverse problem [9], and accounting for the confounding effects of the presence of noise in experimental displacement data. Regularization is needed to address ill-posedness and noise, but this technique is accompanied with its own challenges such as determining appropriate regularization parameters. Ultimately, synthetic test cases are required to ensure inverse model accuracy, but design of appropriate tests requires proper modeling of experimental noise and avoiding the inverse crime [10].

Most current TFM analyses maintain two key assumptions of the hydrogel: 1) the modulus value remains unchanged and is spatially homogeneous [5–8, 11–13], and 2) it is incompressible [7, 12]. Regarding the first assumption, we have previously developed an approach to estimate changes in the local spatial distribution of the hydrogel moduli [14]. This approach has demonstrated that cells actively degrade and stiffen local hydrogel moduli, the latter induced in part by the presence of collagen in the stiffened regions. Regarding the second assumption, we have recently observed significant compressibility in hydrogels in the presence of cellular contraction [4]. To our knowledge this phenomenon has not been investigated in relation to whether it can lead to large errors in inverse-model-predicted mechanical quantities.

In the present study, we developed a robust finite-element inverse-modeling approach to accurately extract local, spatially-varying hydrogel moduli while accurately recovering the observed hydrogel compressibility. As an improvement to our previous approach, we enabled modulus predictions capable of spanning multiple orders of magnitude. This involved use of an exponential relationship for the hydrogel moduli and multiple stages of L-BFGS optimization with progressively lowered regularization parameters, allowing stable recovery of local moduli. We extensively validated our approach by means of synthetic test cases in which the degree of compressibility was the same between ground-truth and inverse-model simulations. We next assessed errors in predicted mechanical quantities arising from improper handling of compressibility through further test cases wherein hydrogel incompressibility was assumed. We then demonstrated our approach by employing novel mitral valve interstitial cell (MVIC) TFM data, wherein we observed orders of magnitude changes in local moduli and identified compressible behaviors in the surrounding poly(ethylene) glycol (PEG) hydrogels.

## 2. NUMERICAL METHODS

### 2.1 Summary of Approach

Our previous experimental observations confirmed the presence of significant cell-contraction-induced compressibility in hydrogels deformed by human induced pluripotent stem cell endothelial progenitors [4]. The Jacobian at referential coordinates in these hydrogels *J* (**x**_0_) = det **F**(**x**_0_), where **F**(**x**_0_) = **I**+∇**u**(**x**_0_) is the deformation gradient tensor, ranged from less than 0.9 to more than 1.1, definitively demonstrating compressible behaviors in the hydrogel. This observation was consistent with known porous behavior of hydrogels, explaining compression with fluid exudation and expansion with fluid swelling. Notably, deviations from *J* = 1 occurred in contractile regions near the cells, zones of great interest to inverse TFM analyses because they are often adjacent to cell surface protrusions where cells exert their largest traction forces [15] and focally deposit ECM material [14]. Accurate determination of cellular force-generation behaviors on these protrusions depends on accurate material modeling of the surrounding hydrogel. Contrary to our recent observations, however, previous inverse analyses to determine these behaviors, including our own [14] and others [12], assumed that the hydrogels were incompressible. The effects of this assumption have not yet been assessed. Our previous inverse modeling approach found that heterogeneous hydrogel moduli near porcine aortic VICs ranged 30 Pa – 250 Pa. A combination of improperly assuming hydrogel incompressibility and the cells’ low metabolic activity, however, may have biased the predicted modulus range to be smaller than the true modulus range of more-active cells.

Based on these findings, we developed a novel finite-element inverse-modeling approach for 3D TFM utilizing a nonlinear *compressible* constitutive model. Moreover, to more fully account for cell metabolic activities, such as MMP-secretion and deposition of extracellular matrix (ECM) [14], we determined the spatially-varying hydrogel modulus field. Importantly, to extend our previous inverse modeling methods to predict hydrogel modulus modification that could only detect modest changes, our new improved approach allowed detecting changes spanning multiple orders of magnitude. Our approach also utilized a variety of numerically-stabilizing features.

We evaluated the accuracy and robustness of the inverse model to make predictions in extensive synthetic test case studies. Importantly, for the first time we demonstrated the need for properly accounting for compressibility in large-deformation 3D TFM.

### 2.2 Inverse Finite Element Model

#### 2.2.1. Hyperelastic Compressible Constitutive Model

We modeled the hydrogel as a neo-Hookean nonlinear hyperelastic compressible solid and assigned the form from [16] with a spatially-varying material parameter

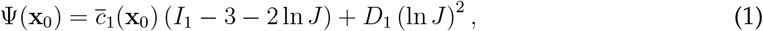

where *I*_1_ = tr **C**, the full first invariant of the right Cauchy-Green deformation tensor **C** = **F**^*T*^ **F**. 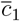 corresponds to one half of the shear modulus and *D*_1_ corresponds to one half of the first Lamé parameter in the infinitesimal strain limit. This approach allows for modulation of the level of compressibility by the value of *D*_1_, with lower values corresponding to increased compressibility. In contrast, this formulation is also applied to approximate incompressibility by adopting *D*_1_ *>> c*_1_, known as “near-incompressibility” [14].

While we have previously utilized a similar form for the material model in [14], the range of 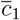 was limited. However, very large moduli of some ECM components such as *de novo* collagen in isolation have been reported to be ≥ 1 GPa [17], suggesting a much wider range is required to capture actual local hydrogel mechanical behaviors. In addition, ECM depositions as well as zones of enzymatic degradation in TFM studies tend to be focal, indicating the presence of large spatial gradients in 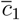. To accommodate these important TFM features we redefined 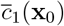 as

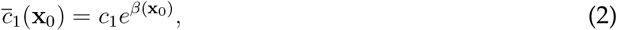

where *c*_1_ = 54 Pa was the “far-field” or unmodified value based on rheometric measurements of the hydrogel shear modulus *µ*; *c*_1_ corresponds to 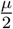 in the infinitesimal strain limit [14]. *β*(**x**_0_) was a scalar variable field defined in the hydrogel volume. With this form, *β <* 0 is identified with local degradation of the hydrogel, *β* = 0 is no modification, and *β >* 0 is local stiffening. This exponential form provided numerical benefits when the degrees of freedom of *β* served as control variables in the inverse model. The exponential relationship did not permit negative modulus because *e*^*β*^ *>* 0 for all values −∞ *< β <* ∞, thus granting stability for small modulus. Gradient computation was also numerically stabilized by the exponential formulation for large local updates in modulus between iterations in optimization. For example, if 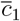 needed to change locally from 54 Pa to 54 kPa, *β* needed to only change from *β* = 0 to ≈ 7. Our previous approach utilized a linear representation, which in this example would have needed to change by 3 orders of magnitude, causing numerical instability that our new approach remedied.

#### 2.2.2 Optimization and Regularization

Building on the above, we developed a finite-element inverse-modeling framework to find *β*(**x**_0_) such that the corresponding simulated hydrogel displacement field **u**_*sim*_ matched the target hydrogel displacement field **u**_*tar*_ (Figure 1). This was done by minimizing the following objective functional with respect to *β*_*sim*_,

**Figure 1.**
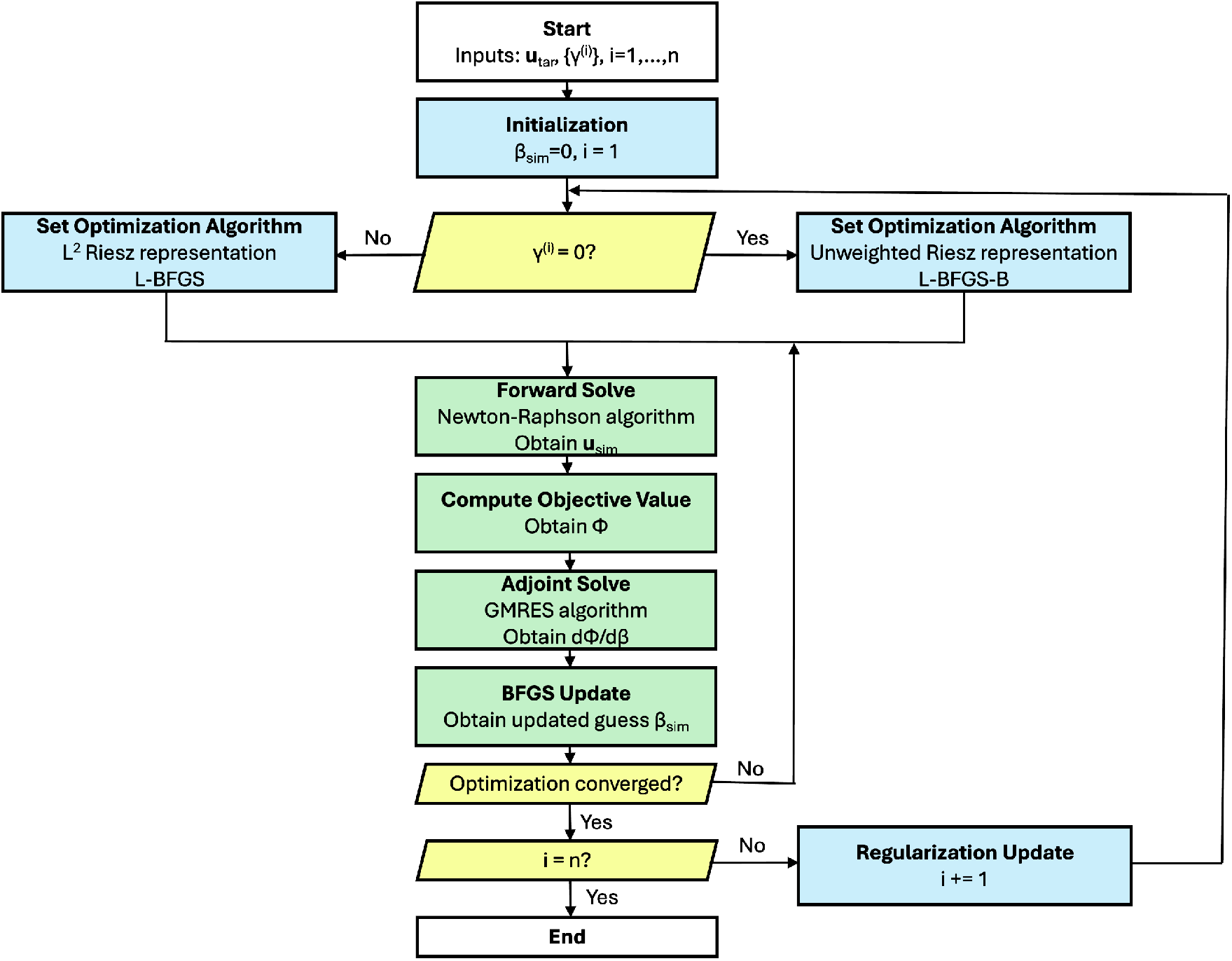
Flowchart of the inverse model. Starting from the target displacement field on the hydrogel and a schedule of regularization parameters, this flowchart demonstrates the components of the inverse-model pipeline, including multi-stage regularization parameter updates. White boxes denote start and end, blue boxes specify updates to configuration, yellow rhombi indicate conditionals, and green boxes are computation steps.

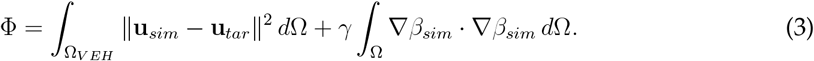

In practice, the second term on the right hand side (a form of Ridge or Tikhonov regularization) was required to stabilize the solution. Here, *γ* was a regularization parameter obtained using realistic test case simulations, Ω was the volume of interest, and Ω_*VEH*_ was the “volume within the event horizon,” which we now define.

Typical TFM micro-sphere displacement magnitudes tend towards zero as the distance from the cell surface increases. Given the associated noise in the experimental data, this suggests a maximum distance from the cell surface beyond which displacements cannot be reliably determined. We thus introduced a “Volume within the Event Horizon” (Ω_*VEH*_), where by “event horizon” we refer to the boundary threshold where micro-sphere displacement magnitudes could be reliably determined. The threshold value was determined by computing the 0-centered moment of micro-sphere displacement magnitudes far from an exemplar cell, *σ*^2^, and then we chose the value 3*σ* to encompass a large extent of noise. Next, the volume within the event horizon for each cell was determined by including tetrahedral elements in hydrogel that had any adjacent nodal displacement magnitudes above the threshold value. We restricted the displacement-matching term of the objective to the volume within the event horizon and also fixed degrees of freedom of the control variable *β*_*sim*_ outside it to 0. This dimensionality reduction simplified the optimization problem by accelerating convergence compared to working with the full problem volume.

#### 2.2.3 Numerical Implementation with L-BFGS

Optimization was performed by bounded and unbounded versions of the L-BFGS (Limited-memory Broyden-Fletcher-Goldfarb-Shanno) algorithm [18], which required functional evaluations Φ(*β*_*sim*_) and gradients of the functional. Functional evaluation at each iteration involved forward-model solves to obtain displacements corresponding to 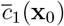 encoded by the current *β*_*sim*_. Gradients were computed by solving the adjoint problem with respect to a particular inner product, as we will describe below. Convergence was detected by either the objective functional value falling below the tolerance 10^−8^ or the gradient magnitude falling below 10^−6^. Finiteelement forward solves were performed in `FEniCS` [19–26] while `FEniCS-adjoint` [27] automatically assembled the adjoint problem and computed gradients.

In addition to the exponential representation of *c*_1_ and regularization methods described above, we adopted a multi-stage regularization scheme for further stabilization. This scheme addressed instability resulting from the initial guess (*β*_*init*_ = 0) being far from the final values. In that case, the first iteration could introduce extreme spatial variations in moduli. Subsequently, the forward model would solve for simulated displacements starting far from equilibrium, which can be numerically unstable without extensive load stepping [12]. To penalize the high modulus gradients early in the optimization process that caused this instability, we initialized the regularization parameter to an appropriately large value and progressively decreased it in multiple stages. For example, a schedule of values input to the model may have been *γ*^(1)^ = 10, *γ*^(2)^ = 0.3. The specific parameter values were determined by calibration to realistic test cases according to the noise level and each cell’s numerical stability requirements. Optimization was run to convergence at each stage and then restarted with the next value. If the final regularization parameter value was *γ* = 0, we utilized the bounded variant of the algorithm, L-BFGS-B, implemented in `scipy` [28] to stabilize modulus gradients in lieu of the regularization penalty. Otherwise, we utilized the unbounded version of L-BFGS implemented in `MOOLA` [29] with line search according to Strong Wolfe conditions [30]. Whereas our previous approach applied the bounded version of L-BFGS to obtain stability over a limited modulus range with a constant regularization parameter, our new multi-stage approach was designed for stabilized recovery over a much larger range.

#### 2.2.4 Forward Simulations with Finite Element Method

Within each iteration of L-BFGS, in order to compute the current objective-functional value, it was first necessary to compute simulated displacements corresponding to the current guess of 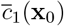. These simulated displacements were the solution of the nonlinear PDE system corresponding to minimization of total hydrogel strain energy

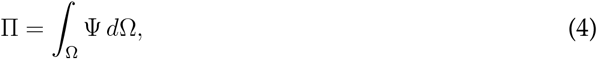

where Ψ was the strain energy density of our hyperelastic compressible constitutive law (Equation 1). We prescribed Dirichlet boundary conditions on both the cellular surface and the outer, surrounding box according to the target displacement field restricted to those surfaces. This boundary condition both forced simulated displacements to match the target at the cellular surface and accounted for hydrogel displacements propagating to the edge of the volume of interest. We discretized the solution displacements in the first-order Lagrange basis on the tetrahedral mesh of the hydrogel. In accordance with standard finite-element procedures, we solved for degrees of freedom satisfying the first-order optimality condition by the Newton-Raphson method. Newton-Raphson iteration progressively updated **u**_*sim*_ such that the residual was minimized; the *j*th iterate 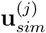 was updated according to

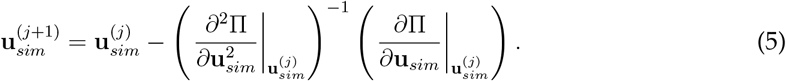

Linear solves represented by the last term were accomplished by GMRES preconditioned with AMG, as implemented by `PETSc` [31] and `Hypre` [32] respectively, with the software’s default tolerance criterion for Krylov solvers [33]. The nonlinear system was assembled by `FEniCS` with third-order quadrature and we recomputed the consistent tangent operator 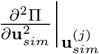 at each Newton-Raphson iteration [34]. We chose to repeatedly reassemble this operator, rather than only assemble the operator once in a quasi-Newton scheme, because we expected substantial non-linear material behavior that would vary both spatially and between L-BFGS iterations. Computations were performed with parallelization facilitated by `OpenMPI` [35].

We halted the Newton-Raphson iteration either upon convergence of the residual to 0 or upon reaching a maximum number of iterations. Convergence was detected by an absolute residual magnitude of *<* 10^−10^ or a relative magnitude *<* 10^−9^. We limited the iteration count because we recognized that forward solves were performed in an inverse-modeling context, thus forward solves in the middle of inverse optimization were approximations and did not need to achieve full accuracy. The specific maximum number of iterations, 8, was calibrated to a synthetic test case. We also leveraged the outer inverse modeling loop by passing previously-computed displacement fields as initial guesses to the next forward solves between inverse model iterations, as in [12]. Successive forward solves between line search and L-BFGS iterations were for updating displacements according to progressively updated modulus states, thus we suspected the difference between successive simulated displacement fields would become smaller as optimization progressed. Reduced initial residuals would then accelerate convergence of forward solves. With the solution of each forward simulation, we computed the current value of the objective functional (Equation 3) for L-BFGS. It remained to compute the current gradient of the objective functional with respect to the control variables *β*_*sim*_, which required an additional step.

#### 2.2.5 Gradient Computation with Adjoint Method and Riesz Representer

As a gradient-based optimization algorithm, L-BFGS required the gradient of the objective with respect to *β*_*sim*_ to perform its optimization. The objective Φ depended on *β*_*sim*_ both directly in the regularization term and indirectly in the displacement matching term through **u**_*sim*_ (Equation 3]). We previously described the relationship between simulated displacement and the current guess *β*_*sim*_ through the nonlinear PDE (Section 2.2.4). To differentiate through this relationship and assemble the full derivative of the objective, 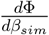, we deployed the adjoint method as in our previous approach [14]. Additionally, we weighted the gradient of the objective over space to equalize the influence of each degree of freedom of *β*_*sim*_, as we will describe in detail below. This weighting improved numerical stability over our previous approach.

`FEniCS-adjoint` automatically determined and solved the appropriate adjoint problem, which we briefly describe here (see [27] for a complete derivation). We sought the total derivative of the objective functional, 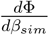, and introduced an adjoint field *ζ* discretized on the tetrahedral hydrogel mesh with the first order Lagrange basis. The adjoint problem to solve for *ζ* was linear and given by

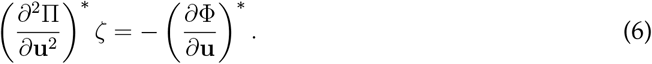

After solving the linear system, we assembled the full derivative by

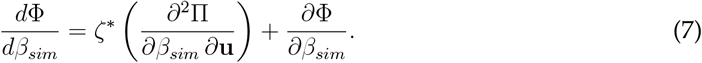

Note that we have been careful to refer to this object as a *derivative*, not a gradient, as yet in this presentation of the adjoint finite-element technique. The derivative is a linear functional, an element in the dual of a vector space. A gradient is a particular vector: a “Riesz representer” corresponding to the derivative that is only defined with respect to a particular inner product in a Hilbert space [36]. L-BFGS specifically requires a gradient, not a derivative, because it is added to vectors in the same space. Most software elides this distinction, including our previous inverse-modeling approach, by treating input vectors out-of-context and thus implicitly assumes an inner product that gives all components the same weight. In our finite-element context, however, we had geometrically-meaningful choices of inner product, such as that corresponding to *L*^2^(Ω), which implied a different gradient than the unweighted default choice. The software package `MOOLA` [29] implemented L-BFGS with a `FEniCS-adjoint` interface for exactly this purpose, thus we incorporated it into our new approach.

We chose the *L*^2^(Ω)-inner-product to weight updates of modulus according to the volume influenced by each *β*_*sim*_ degree of freedom. This weighting smoothed the predicted spatial gradients of modulus and thus stabilized forward solves compared to our previous unweighted approach. Numerically, the components of the gradient 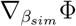 in the Lagrange basis were related to components of the derivative 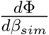 in the dual Lagrange basis by the mass matrix *M*,

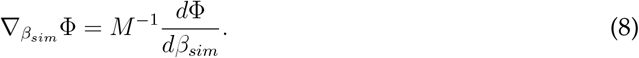

Additional modifications to L-BFGS were implemented by `MOOLA` to perform computations with the compatible norm, we refer the reader to [37] for the details.

#### 2.2.6 Post-Processing

The inverse model’s output was the final modulus field prediction 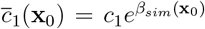. The simulated displacements corresponding to this modulus, **u**_*sim*_, were the solution to the forward model with the optimal *β*_*sim*_. We computed simulated *J* from simulated displacements *via* standard finite-element shape-function routines. We evaluated the match between simulated and target *J* fields as it was a necessary part of adequately accounting for compressibility.

When computing traction forces, we carefully chose how we discretized them over the cellular surface. With our exponential representation of *c*_1_, traction forces were simulated as living in an exotic space that was discontinuous across triangular surface elements and varied exponentially in the interior of the triangles. We therefore projected surface traction forces onto a vector first-order Lagrange basis with nodal values at the cellular surface. This choice of basis had the same dimension as the number of degrees of freedom for *β*_*sim*_ located on the cellular surface and thus provided a reasonable approximation of corresponding traction. Formally, traction vectors are located in the current configuration and are also oriented according to the current configuration. Thus, analogues of traction force in the referential configuration can be misleading, therefore we utilized the current configuration for our traction analysis. `FEniCS` natively performed computations only in the reference configuration, so we utilized the first Piola-Kirchoff stress tensor **P** to compute traction forces in the current configuration. Specifically, **P** is a two-point tensor that maps normal vectors in the referential configuration **n**_0_ to traction vectors in the current configuration per area in the referential configuration using

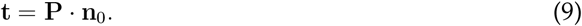

Next, we evaluated the spatial distribution of strain energy as it captured the contractile behaviors of the embedded cells. The kinematic quantities *I*_1_ and *J* included in its definition (Equation 1) were element-based. As with traction, however, the exponential dependence on *β* made strain energy density vary exponentially within elements. We thus projected it into an element-wise space in order to control intra-element variation while respecting its natural interpretation as a per-element quantity in the absence of hydrogel modification. We computed total strain energy in the volume of interest, on the other hand, by `FEniCS`’s builtin integration routines with the full precision version of the strain energy density field living in the exotic, exponential space.

### 2.3 Synthetic Test Cases

#### 2.3.1 Test Problem Design

To thoroughly validate inverse-model predictions of 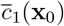, total hydrogel strain energies, and net traction forces, we generated several test cases with realistic cell geometries and prescribed 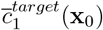 with characteristics reflective of real cell-modified hydrogel. We generated target displacements from forward finite-element simulations with boundary conditions from real experimental displacement data. Based on previous findings of porcine aortic valve interstitial cell deposition patterns [14], we prescribed two stiffened regions near contractile regions of the cellular surface and degradation elsewhere. To represent potentially very large modulus (as might occur in *de novo* synthesized collagen), we prescribed a maximum thousand-fold increase in modulus in stiffened regions. We set the minimum modulus to one third the far-field value, a moderate amount in accordance with the association of compliant regions with hydrogel degradation [14]. The degree of compressibility was set to fit real cell-modified hydrogel behavior. We performed multiple tests with this test problem (Table 1) to demonstrate 1) inverse model validity when properly accounting for compressibility and 2) improperly accounting for compressibility causes the inverse model to produce inaccurate predictions. For all test cases, we compared 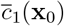 fields between target and inverse-model output. High quality agreement of modulus was of primary interest due to its interpretation as a metric of cell secretion and deposition activity. We additionally compared quantities of interest including displacements, *J*, traction forces, and strain-energy densities.

**Table 1.** Test case descriptions. Parameters for each of the four cases. Mesh length refers to the characteristic length scale applied to generate the tetrahedral mesh of the hydrogel domain. 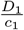 values indicate the material parameter settings in ground-truth and inverse-model forward simulations; smaller values are more compressible. “Exact” target displacements indicate the output of the forward model was directly input to the inverse model and “Noise Added” indicates synthetic noisy micro-sphere displacements were interpolated to generate the target displacements. The first two cases established the validity of the approach. The last two cases demonstrated the effects of improperly assuming incompressibility.

| Ground-Truth<br>Mesh Length | Ground-Truth $\frac{D_1}{c_1}$ | Target<br>Displacements | Inverse-Model<br>Mesh Length | Inverse-Model $\frac{D_1}{c_1}$ |
| --- | --- | --- | --- | --- |
| 2.86 $\mu\text{m}$ | 1 | Exact | 2.86 $\mu\text{m}$ | 1 |
| 2 $\mu\text{m}$ | 1 | Noise Added | 2.86 $\mu\text{m}$ | 1 |
| 2 $\mu\text{m}$ | 1 | Noise Added | 2.86 $\mu\text{m}$ | 10 |
| 2 $\mu\text{m}$ | 1 | Noise Added | 2.86 $\mu\text{m}$ | 100 |

#### 2.3.2 Validation of Modulus and Traction Forces

First, we validated the accuracy of the inverse model in recovering modulus while using the same degree of compressibility as ground truth in a scenario with perfect target displacement information. This test was for assessing *numerical* validity; we expected the inverse model to recover the target field nearly perfectly. Success would show that the inverse model could detect the true modulus field and navigate to it starting from guess *β*_*init*_ = 0. While generating target displacements, we used the same mesh discretization as the inverse model for maximum compatibility.

Next, we established that the inverse model could adequately recover modulus when the target displacement field was obtained from Gaussian-process interpolation of noisy micro-sphere displacement data, as is the case in real applications. We generated clean target displacements **u**_*tar*_ from a forward simulation with a finer-resolution version of the hydrogel mesh (characteristic length 2 *µ*m) to avoid the inverse crime [10]. At the real micro-sphere positions corresponding to the real cell, we evaluated the clean target displacement field to obtain ground-truth synthetic micro-sphere displacements. We sampled noise from a Gaussian distribution for each position according to the fit from real far-field micro-sphere data. We then added the noise to the synthetic ground-truth micro-sphere displacements to create synthetic noisy micro-sphere displacement data. We fit a Gaussian Process Regression (GPR) model to this synthetic micro-sphere data and interpolated it onto the standard version of the hydrogel mesh with characteristic length 2.86 *µ*m. We provided the resulting noisy displacement field to the inverse model and denoted it **u**_*tar*_ + *δ*. We set the same degree of compressibility in the inverse model as in the ground-truth simulation to validate the case in which hydrogel compressibility is properly handled.

#### 2.3.3 Effects of Compressibility on the Inverse Model

We next evaluated the effect of *improperly* accounting for compressibility in our inverse-modeling approach. In our material model formulation (Equation 1), 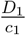 is larger the more incompressible a material is. To determine whether accuracy is substantially deteriorated when assuming too-little compressibility, such as if we were to utilize a previous version of the inverse model pipeline [14], we increased the ratio 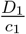 in the inverse model to two different values, 10 and 100, both larger than the ground-truth ratio. These ratios approximated incompressibility through a nearly-incompressible formulation. We didn’t assign any larger ratios as they would induce substantial volume locking in our forward solver [34]. We input the same noisy target displacements **u**_*tar*_ + *δ* to the inverse model as in the previous test. We expected poor recovery of the target modulus field due to improperly accounting for compressibility.

## 3. NUMERICAL RESULTS

### 3.1 Validation of Modulus and Traction Forces

#### 3.1.1 Exact Target Displacements

First, we evaluated the test case in which exact ground-truth displacements were input to the inverse model. This test was performed to assess numerical validity of our approach’s recovery of the target modulus field. We assigned the same amount of compressibility in both the ground-truth simulation and in the inverse model, 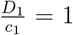, to establish accurate recovery when compressibility is properly modeled. The far-field displacement cutoff was set to 0.38 *µ*m in accordance with the noise level in real data. We found that assigning only 2 MPI processes to the inverse-model program was sufficient to improve runtime before process synchronization time began to degrade performance. Additionally, we set the maximum number of Newton-Raphson iterations to 8 because solves taking more iterations than that stalled in convergence.

Regularization in this context only served a numerically-stabilizing role, thus by trial-and-error we searched for an adequate regularization parameter schedule that ended with no regularization. The starting parameter *γ*^(1)^ = 10 was the smallest value we found for which the first stage converged. We also found that subsequent parameter values could not decrease too quickly lest the model diverge, thus we prescribed two more stages *γ*^(2)^ = 0.001, *γ*^(3)^ = 0. In the first two stages, the choice of *L*^2^ Riesz representation was necessary to smooth intermediate modulus guesses so that forward solves could converge without load stepping. At the final stage, with no regularization, we found that using the unweighted vector Riesz representation and the bounded version of L-BFGS (uniformly −2 ≤ *β* ≤ 20) was more effective for capturing small local details in modulus than the unbounded version with *L*^2^ Riesz representation. This was likely the result of using the exact target displacements and the same mesh discretization as ground-truth simulation; in this case, during the final stage, the predicted 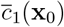 required few improvements in regularity to promote forward-model convergence.

The inverse model converged after its three stages and output a predicted modulus with excellent agreement with the target (Figure 2(a),(b)); the average relative error in modulus in the volume of interest was 1.06%. The final forward solve to produce simulated displacements converged, confirming that our maximum Newton-Raphson iteration count (8) did not deteriorate the quality of the final inverse-model prediction. The maximum nodal error between target and simulated displacements was 0.043 *µ*m. For reference, the maximum nodal difference between the target displacement field and displacements corresponding to an unmodified, uniform modulus field was 3.053 *µ*m. These studies established that our approach was able to converge to the correct modulus field by improving its displacement predictions.

**Figure 2.**
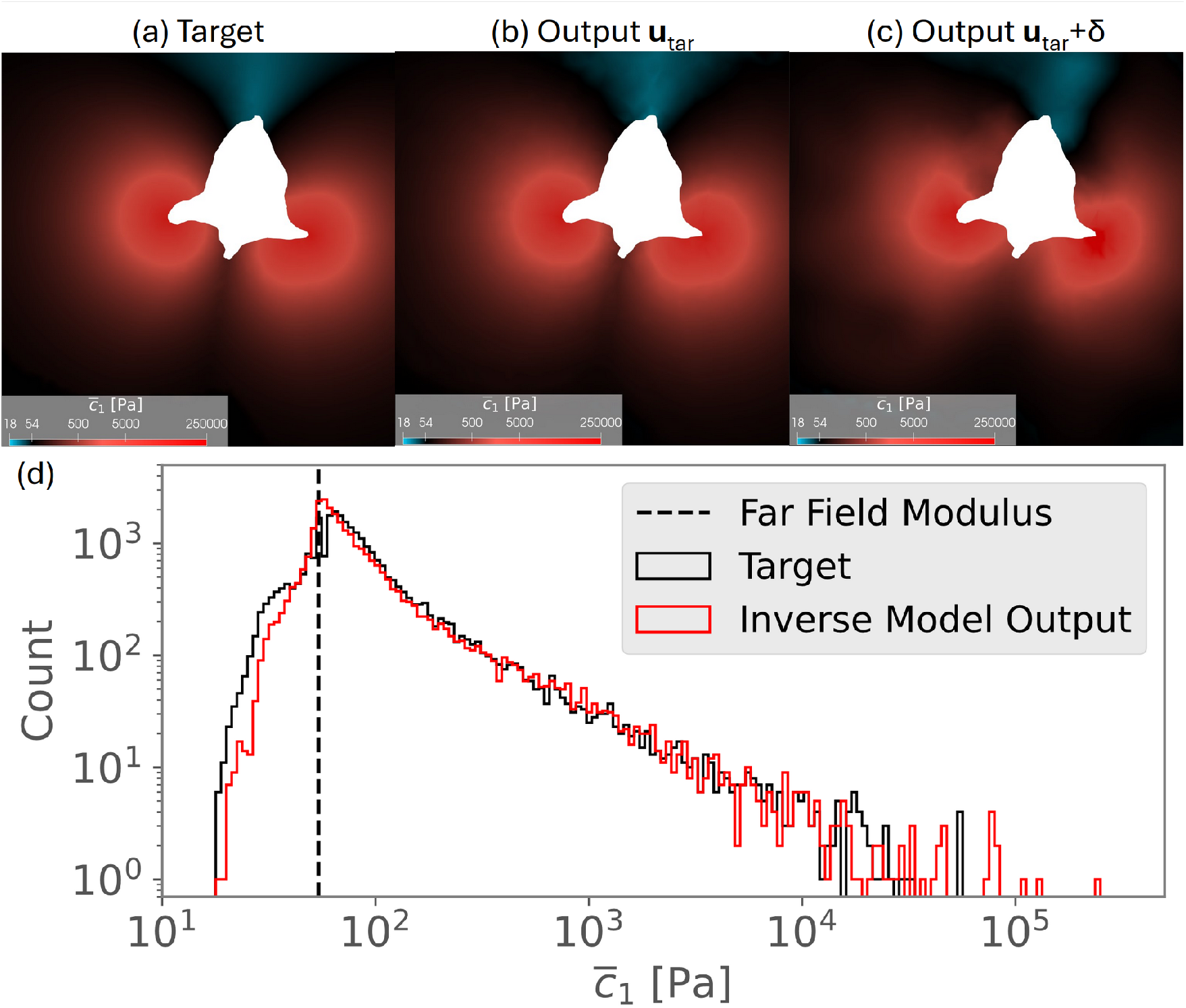
Comparison between prescribed target modulus field and inverse-model output modulus. Top row is a representative cross-section of hydrogel modulus. (a) The prescribed target modulus field on the refined version of the mesh utilized to generate target displacements. (b) The inverse-model output for the case of exact, no-noise target displacements **u**_*tar*_ where ground-truth simulation and inverse model share the same mesh. (c) The inverse-model output for the case of target displacements with noise added **u**_*tar*_ + *δ* and different meshes between ground-truth simulation and inverse model. (d) Comparison of distributions of nodal target modulus values (interpolated onto the mesh that was employed by the inverse model) vs. nodal inverse-model output modulus values for the case with noisy displacement inputs. Only nodal values within the event horizon are included in the histogram because outside values are fixed to the far field value 54 Pa. 3 outlier nodal values with large modulus 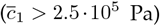 of total 28,171 are excluded.

#### 3.1.2 Effects of Noise

Next, we input the target displacement field with a realistic distribution of noise to the inverse model. Whereas the previous test established both the necessity and effectiveness of numerically-stabilizing components of our approach in ideal conditions, herein we evaluated effectiveness under realistic conditions. Again we assigned the same amount of compressibility, 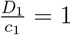, in both the ground-truth simulation and in the inverse model. The far-field cutoff value was 0.38 *µ*m as before. We searched for the regularization parameter schedule that would admit the best match between target and simulated modulus fields. A single stage of regularization with value *γ*^(1)^ = 0.3 was optimal. The inverse model converged quickly, taking only 1 hour 6 minutes on a desktop with a 13th Generation Intel i9 processor and 125 GB of allocated RAM. The final forward model to provide simulated displacements converged without early stopping, verifying simulated displacements appropriately corresponded to the predicted 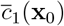.

The output modulus field matched the target well, though with some degraded quality compared to the previous case with perfect displacement information (Figure 2). The average relative error in modulus in the volume of interest was 17.2%. The total strain energy and volumes of stiffened and degraded regions were also recovered well (Table 2). We compared the noisy displacement field with the inverse model’s simulated displacement field along with corresponding *J* values (Figure 3). The maximum nodal difference between the displacements was 0.466 *µ*m, close to the event noise level cutoff value of 0.38 *µ*m. Likewise, the average difference in elementwise *J* was 0.006. For reference, the average difference between clean target *J* (computed using the inverse model’s mesh discretization) and noisy target *J* was 0.009, indicating that the inverse model was able to recover local expansion and contraction within the margin of error.

**Table 2.** Total strain energy, stiffened volumes, and degraded volumes of ground-truth target simulation vs. inverse-model prediction with various levels of compressibility. Target displacements provided to the inverse model in all three cases contained a realistic level of noise. Strain energy was computed within the event horizon by 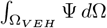, where strain energy density takes the form given in Equation 1. Energy was over-estimated if the hydrogel was modeled as insufficiently compressible, *i*.*e*. the 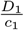 ratio was too large. The stiffened volume was computed according to where predicted modulus 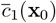 was stiffened by 10-fold over the far-field value *c*_1_ = 54 Pa 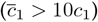, while the degraded volume was computed according to the threshold 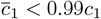.

| | Target<br>$\frac{D_1}{c_1} = 1$ | Simulated<br>$\frac{D_1}{c_1} = 1$ | Simulated<br>$\frac{D_1}{c_1} = 10$ | Simulated<br>$\frac{D_1}{c_1} = 100$ |
| --- | --- | --- | --- | --- |
| Energy [pJ] | 6.78 | 5.30 | 34.8 | 334 |
| Stiffened Vol. [$10^4 \mu\text{m}^3$] | 5.10 | 5.11 | 31.9 | 86.4 |
| Degraded Vol. [$10^4 \mu\text{m}^3$] | 22.6 | 21.5 | 7.62 | 9.20 |

**Figure 3.**
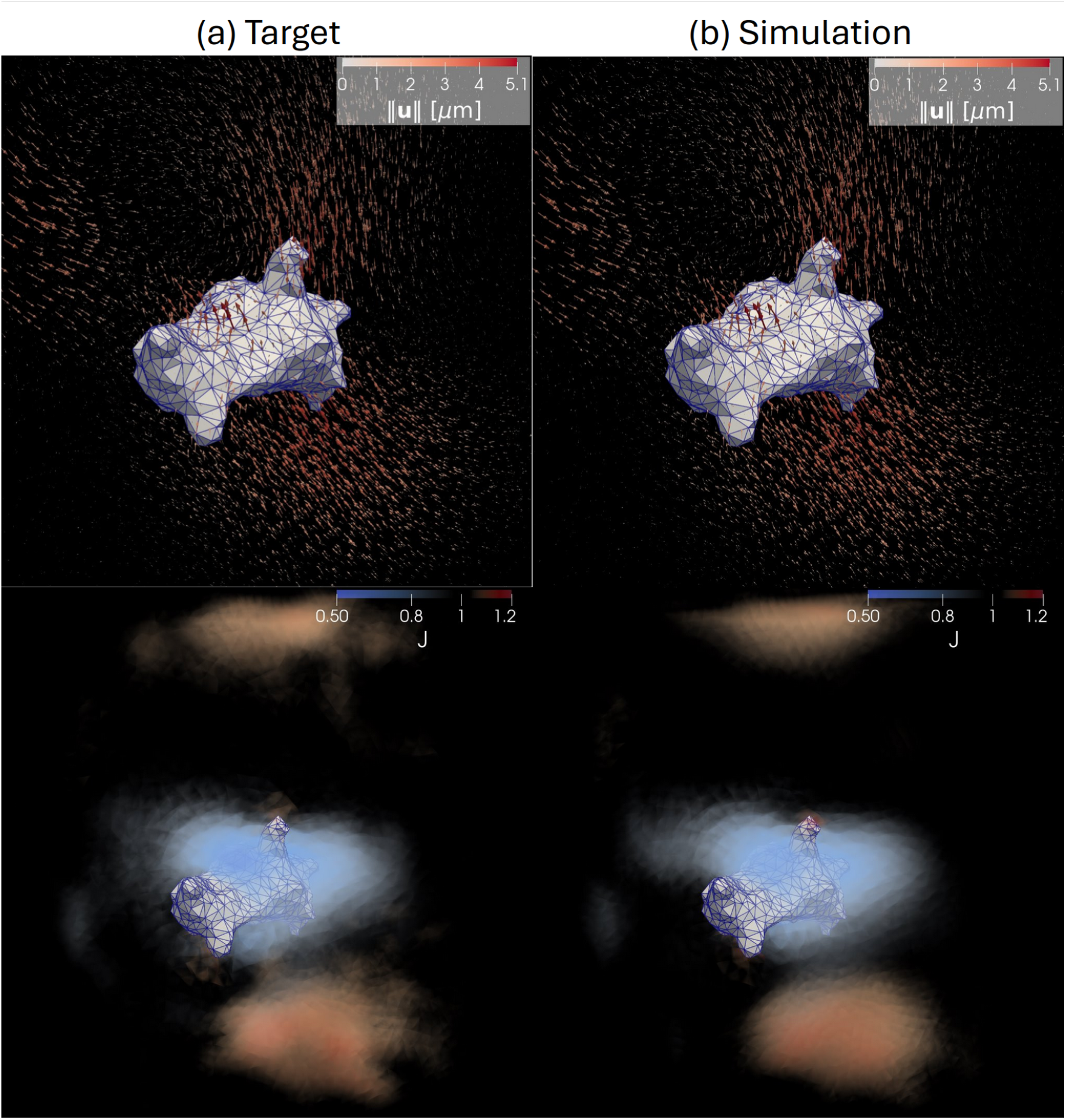
Recovery of kinematic quantities by the inverse model for the test problem with noise. Gray mesh is the cellular surface. The top row is in the current configuration, shaded arrows are displacements, and the bottom row is in the referential configuration. Shaded and transparent elements depict *J* values. (a) The target displacements with noise added **u**_*tar*_ + *δ* supplied to inverse model and the corresponding *J* field. The standard deviation of *J* in the event horizon was 0.039. (b) The inverse-model prediction of displacements and *J*. The standard deviation of *J* in the event horizon was 0.038.

For deeper investigation of inverse-model fidelity, we next computed traction forces corresponding to the inverse-model output. We found that magnitudes of inward and outward traction forces on the cellular surface matched well between the ground-truth simulation and inverse-model prediction (Figure 4). We restricted our analysis to the median 95% of nodal values on the cellular surface due to the presence of outlier values. The root mean squared error in the median traction forces was 29 Pa. In combination with the excellent recovery of spatially-varying moduli across 5 orders of magnitude, our inverse model adequately captured mechanical quantities of interest under realistic noise conditions when compressibility was correctly modeled.

**Figure 4.**
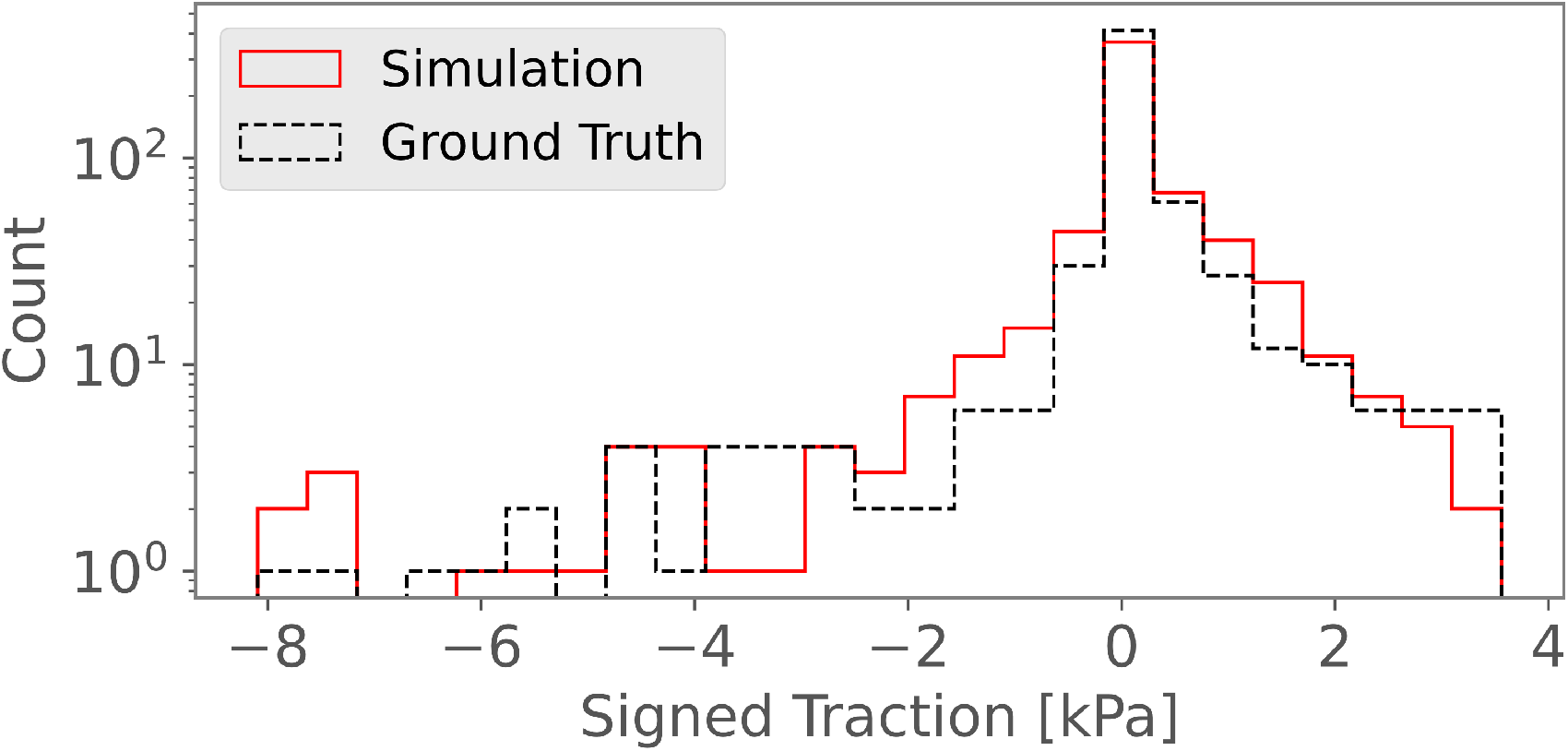
Recovery of cellular-surface traction forces by the inverse model for the test problem with noise. Histogram of the median 95% of signed nodal traction force magnitudes. Positive values are outward-pointed traction forces while negative values indicate inward forces. Ground-truth traction forces were obtained by forward simulation with target modulus field and ground-truth boundary conditions. Simulated traction values were obtained by the inverse-model prediction of modulus and noisy boundary conditions.

### 3.2 Effects of Compressibility on Inverse Model

We next determined the extent of error introduced by the inverse model simulating the hydrogel as incompressible while the ground-truth hydrogel was compressible. Target displacements were obtained from a forward model with the prescribed modulus field (as in Section 3.1.2) and assuming the hydrogel was compressible with 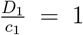. A realistic distribution of noise was added to displacements. These noisy target displacements were input into the inverse model in two different, less-compressible configurations, 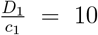 and 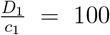, approximating incompressibility through nearly-incompressible formulations as in our previous approach [14]. In both cases, we found it was necessary to exercise two regularization stages, *γ*^(1)^ = 10, *γ*^(2)^ = 0.3. The additional stage over the matching-compressibility case was likely required due to numerical instability arising from the mismatch in ratio between target and simulation. The final regularization parameter value was set to agree with the optimal value found in the previous test with noise. We additionally adopted the matching-compressibility case as a baseline for performance.

The inverse model converged in both cases and we compared predicted quantities of interest against the corresponding ground-truth values. The predicted modulus fields for both under-compressible configurations consistently overestimated the volume and magnitude of stiffening. Predicted degraded zones were shrunk compared to ground-truth, and ground-truth degraded zones were predicted to contain substantial stiffening (Figure 5). The mean relative errors in 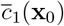 for the 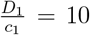 and 100 cases were 415% and 6,650% respectively. Whereas the inverse model with the correct amount of compressibility was able to accurately capture both stiffened and degraded volumes, incompressible material modeling resulted in the inverse model making highly inaccurate predictions of these volumes (Table 2).

**Figure 5.**
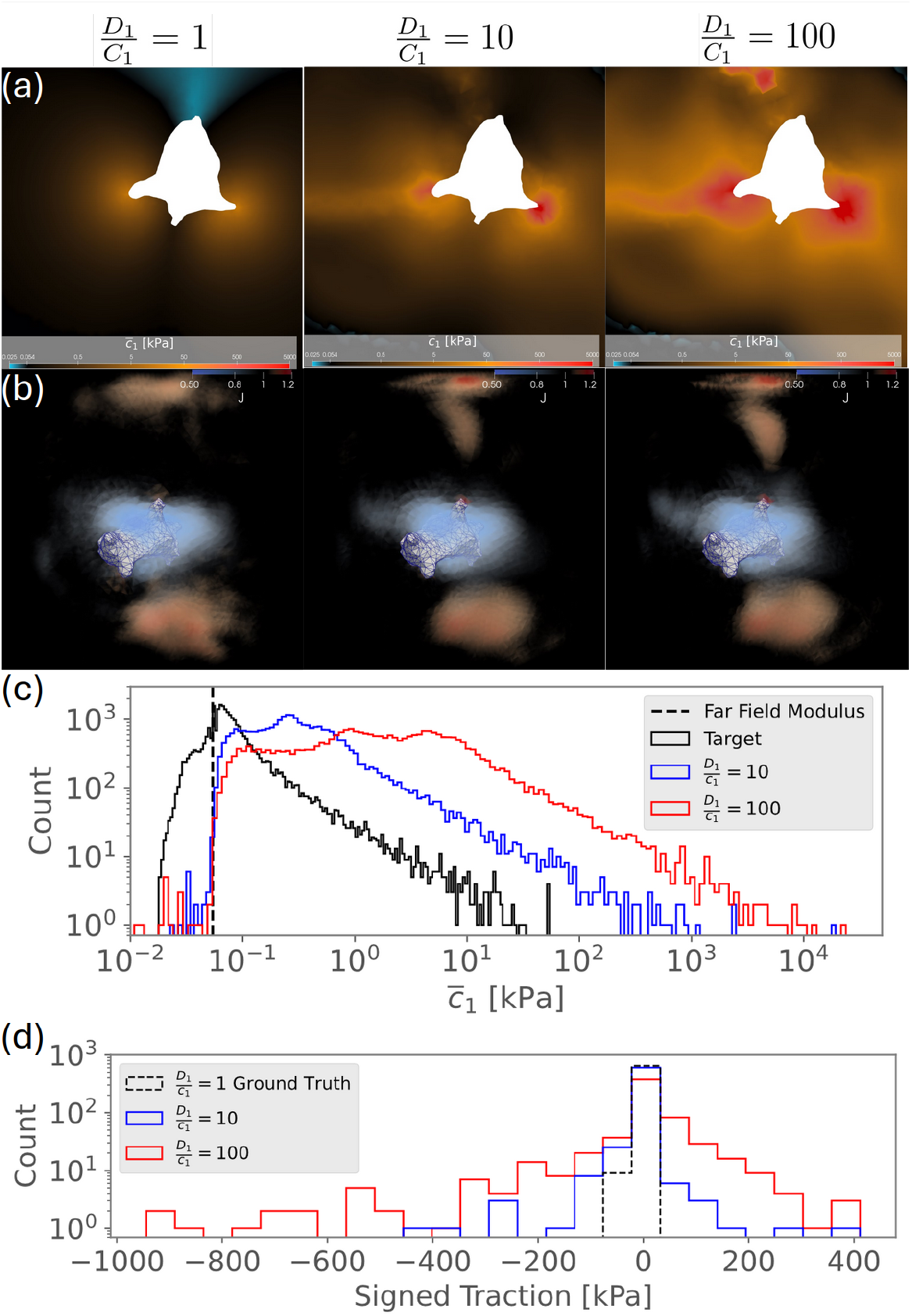
Compressible ground-truth simulation with 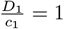 vs. inverse-model simulation with incompressible material models using 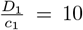 or 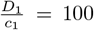. (a) Representative cross-section of hydrogel modulus where the hollow inclusion of the cell is white, unmodified modulus (54 Pa) is black, degraded modulus is blue, the maximum target stiffness is orange, and stiffening greater than the maximum target value is red. (b) Corresponding *J* fields, where transparent elements have *J*≈1, blue is compression *J <* 1, and orange is expansion *J >* 1. Columns left-to-right correspond to: the ground-truth prescribed field and corresponding *J* field from a forward-model simulation with 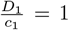, the inverse-model output from target displacements with noise **u**_*tar*_ + *δ* while using 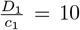 and the corresponding predicted *J*, and the inverse-model output and corresponding *J* using 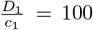. (c) Nodal modulus value distributions within the event horizon where the target field is projected onto the mesh employed by the inverse model. The modulus range is truncated to exclude outliers. (d) Comparison of distributions of the median 95% of nodal signed traction force magnitudes on the cellular surface. Negative values point into the cell while positive values point outward.

The maximum nodal error in displacements for the 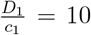 case had magnitude 0.562 *µ*m and the 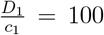 case had magnitude 1.149 *µ*m. These displacement errors were worse than the matching-compressibility case but better than assuming homogeneity, confirming that the inverse model successfully reduced the mismatch between target and simulated displacements. Similarly, the average errors in *J* were 0.008 and 0.01 respectively. These values were close to the difference in *J* arising purely from the presence of noise, 0.009, indicating that the inverse model largely captured the changes in volume in the target, with only a few regions containing large errors in volume ratio (Figure 5). Because the ground-truth displacements and *J* corresponded to a compressible material, however, the 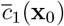 field required to match the changes in volume was far different than the target.

We also assessed the recovery of traction forces and total strain energy for both cases of under-compressible inverse-model configuration. We computed signed traction force magnitudes and found the predicted range of the median 95% traction magnitudes was different from the ground-truth range by multiple orders of magnitude (Figure 5). Root mean squared errors in the median traction forces for the 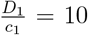 and 100 cases were 76 Pa and 290 Pa respectively, the latter being a 10-fold increase in error over the case of matching compressibility. Total strain energy is a bulk scalar quantity and therefore we expected it to be easier for the inverse model to accurately predict even when compressibility was not matched. Despite this consideration, incompressible modeling resulted in over-estimation of the total strain energy by 5-fold or more (Table 2).

## 4 EXPERIMENTAL DEMONSTRATION

We next demonstrate our methodology using newly-acquired TFM data. The goal here is to demonstrate the utility and insights now possible to extract from TFM experiments with our inverse-modeling approach. Rather than utilizing extant data, we leveraged recent improvements in experimental methodology [4], including higher spatial imaging resolution and doubled micro-sphere densities. These features together yielded a more-detailed picture of the local cellular geometry and induced deformations, providing improved experimental data to evaluate our inverse model capabilities. We further confirmed the presence of compressibility in PEG-hydrogel-based TFM studies to similar levels as we found previously in norbornene-functionalized hyaluronic acid hydrogels [4].

### 4.1 Experimental Data Collection and Processing

Experimental and data processing methods have been presented in detail [3, 14, 38]. In brief, human MVICs were isolated from a consented patient at the Columbia University Biobank for Translational Studies (IRB AAAR6796) and shipped to University of Texas at Austin. Aliquots of the expanded MVICs were stained and integrated into 3D PEG hydrogels prepared with fluorescent micro-sphere density increased to 3 × 10^9^/mL. This improved approach allowed us to utilize micro-spheres at double previous densities and to determine their displacements with higher accuracy. Two MVICs and surrounding micro-spheres were imaged with a laser scanning microscope in the basal state, then treated for 40 minutes with cytochalasin-D, and then reimaged.

We tetrahedralized the imaged hydrogel volume (dimensions 147 *µ*m × 147 *µ*m × 119.23 *µ*m), excluding a 2 *µ*m buffer on all sides to improve the accuracy of interpolated displacements near the outer boundary (Figure 6). All elements were first-order Lagrangian four-node tetrahedra with a 2.86 *µ*m characteristic edge length. The cellular surface mesh was incorporated into the hydrogel mesh as a hollow inclusion after coarsening to have characteristic element side lengths matching the hydrogel mesh using `PyMeshLab` [39, 40]. Mesh generation was performed by `Gmsh` [41] with `netgen` [42] optimization to improve element aspect ratios. We interpolated micro-sphere displacements to nodal points using a GPR model that incorporated an approximation of the variance of the noise in each component of micro-sphere displacements according to the theory described in [43]. Variances were determined by selecting micro-spheres at a distance of 40 *µ*m from the cellular surface (where the displacements were smaller than the voxel size) and computing a 0-centered variance of each component of displacement. The cytochalasin-D-treated state was identified as the referential configuration; actin polymerization was deactivated and the cell was no longer able to generate contractile forces. Element-wise *J* (**x**_0_) values were computed using standard finite-element shape-function routines and analyzed to assess the presence of compressibility.

**Figure 6.**
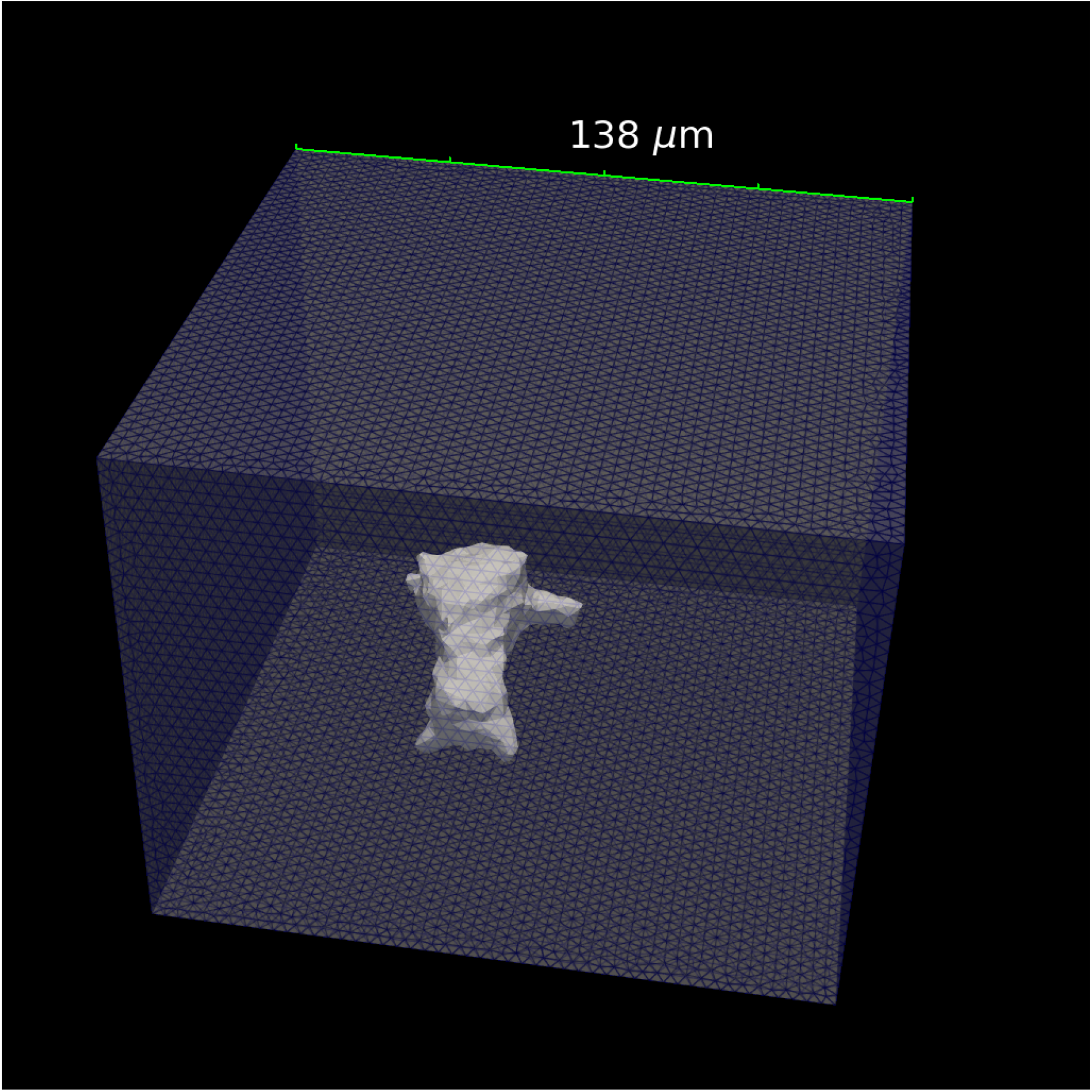
Hydrogel mesh. A tetrahedral finite-element mesh of the hydrogel domain of our simulations with characteristic element side lengths 2.86 *µ*m. The outer surface is semi-transparent with highlighted element edges to delineate structure, and the white mesh is the cellular surface. Dimensions of the box varied according to the size of the accurately-tracked region but were always contained within the imaging volume. The 3D cell geometry was a hollow inclusion.

### 4.2 Experimental Findings

Both MVICs exhibited hydrogel elements where the change in volume exceeded *±*0.05, contrary to the incompressibility assumption that there were no local changes in volume. Such elements were concentrated near the cellular surface (Figure 7). *J* had ranges 0.455 – 1.662 and 0.369 – 1.879 respectively, comparable to the ranges we previously observed, which frequently exceeded 0.9 – 1.1 [4].

**Figure 7.**
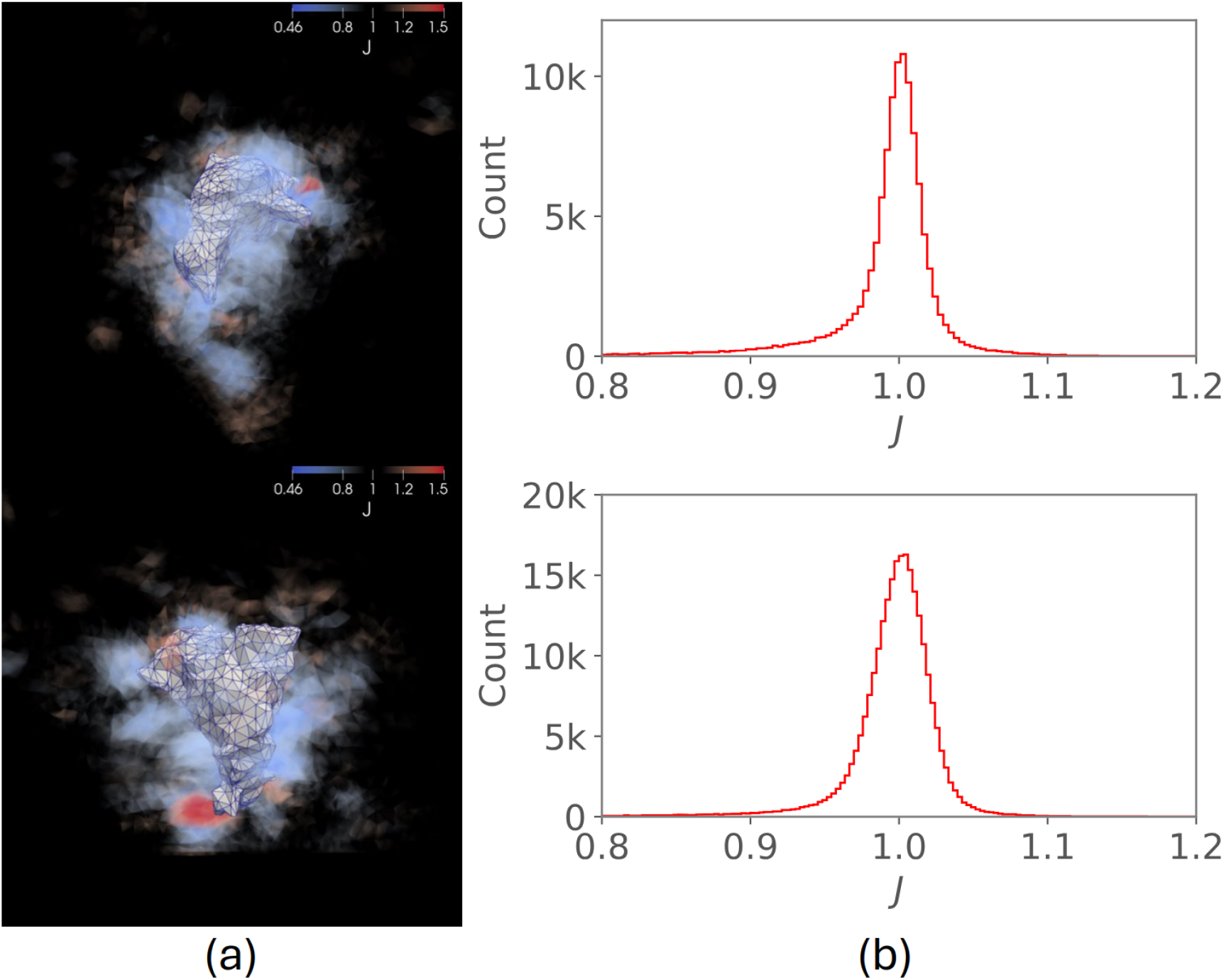
Local *J* values substantially differed from 1 near MVIC surfaces in the PEG hydrogel, violating the traditional incompressibility assumption. (a) Each MVIC surface (gray mesh) in the referential configuration surrounded by a volumetric representation of the *J* field from experimental displacements. Elements are transparent for *J*≈1 and colored blue for compression and red for expansion. Note that the second cell was positioned near the edge of the imaged volume; deviation from *J* = 1 may have continued further away from the cell. (b) Corresponding histograms of *J* values on each element in the hydrogel volume within the event horizon (cutoff value 0.38 *µ*m), range of *J* truncated to highlight representative variation. From top to bottom, means and standard deviations of *J* were 0.992 *±* 0.043 and 0.996 *±* 0.036.

### 4.3 Inverse Model Predictions

First, to determine an adequate level of compressibility 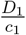, we chose an MVIC and performed numerous forward simulations with varying 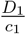. To simplify this initial calibration, we assumed a homogeneous far-field modulus. Resulting *J* distributions were compared between homogeneous simulation and experimental values. We found that 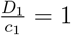 achieved a sufficient fit under the simplifying homogeneity assumption and thus set this level of compressibility in the inverse model. We subsequently justified our choice of this ratio based on inverse-model-predicted *J* fields, as we will exhibit later. We next set a single regularization parameter stage *γ*^(1)^ = 0.3, as was best for the test case. We then processed both of the human MVICs with the inverse model. In the inverse-model output 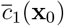 for each cell, a small fraction outlier nodal values were present with extremely large stiffening. Thus, we excluded those nodes from our analysis. The 99.9th percentile smallest and largest moduli in one MVIC were 3.6 Pa and 2.4 MPa, spanning over 5 orders of magnitude (Figure 8).

**Figure 8.**
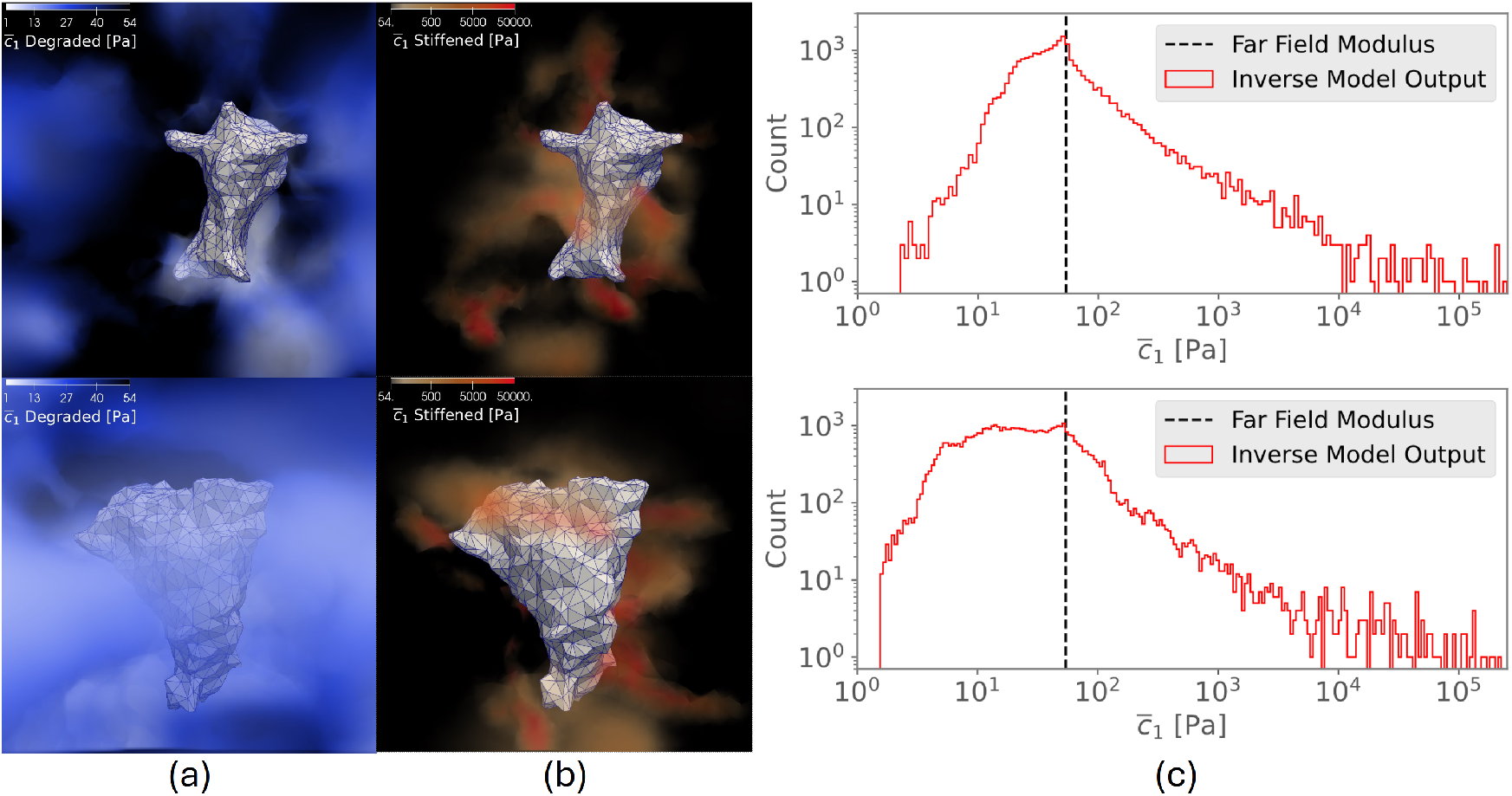
Inverse-model output of hydrogel modulus predictions for each MVIC. For (a-b), MVIC surfaces (gray meshes) are in the reference configuration. Volumetric representations of the modulus field are transparent in unmodified regions with 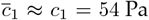 Pa, the far-field value. (a) The degraded region where modulus was below the far-field value. Lighter shades of blue indicate greater compliance and the scale bar is linear. (b) The stiffened region where modulus was greater than the far-field value. Red colors are stiffer and the scale bar is logarithmic to demonstrate intermediate variations in stiffening. The scale is truncated to highlight representative variation without outlier nodal values. (c) Corresponding histograms of nodal modulus values within the event horizon.

We considered nodes where modulus was 10-fold larger than the far-field value, 54 Pa, to be substantially stiffened 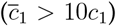, and in this subset the median moduli were 1500 Pa and 1350 Pa respectively. This finding demonstrated that stiffened regions frequently contained points with 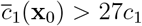, greater than the maximum allowed by our previous approach. We additionally found that accounting for such extreme heterogeneity dramatically altered inverse-model-predicted strain energy in contrast to assuming homogeneity. We ran our forward model with both the inverse-model output 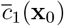 and a uniform material parameter *c*_1_; both cases converged and we computed per-element strain-energy densities. The strain-energy density field for the heterogeneous case often locally differed from the homogeneous case by an order of magnitude or more (Figure 9), which indicated large differences in predicted cellular mechanical behavior when accounting for the full range of heterogeneity. Likewise, the simulated total strain energy in the volume of interest was greater in the heterogeneous case by factors of 27 to 50 (Table 3).

**Table 3.**
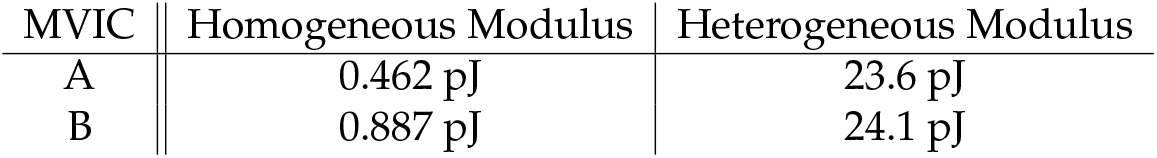
Total cell-contraction-induced strain energies in the hydrogel assuming homogeneous modulus vs. heterogeneous modulus from inverse-model prediction, presented in the order of Figure 7. Strain energy was computed in the volume of interest according to the material model in Equation 1. The homogeneous case assumed unmodified modulus (uniformly the far-field value *c*_1_ = 54 Pa) while the heterogeneous case assigned modulus output 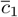 by the inverse model. Values were rounded to 3 decimal digits.

| MVIC | Homogeneous Modulus | Heterogeneous Modulus |
| --- | --- | --- |
| A | 0.462 pJ | 23.6 pJ |
| B | 0.887 pJ | 24.1 pJ |

**Figure 9.**
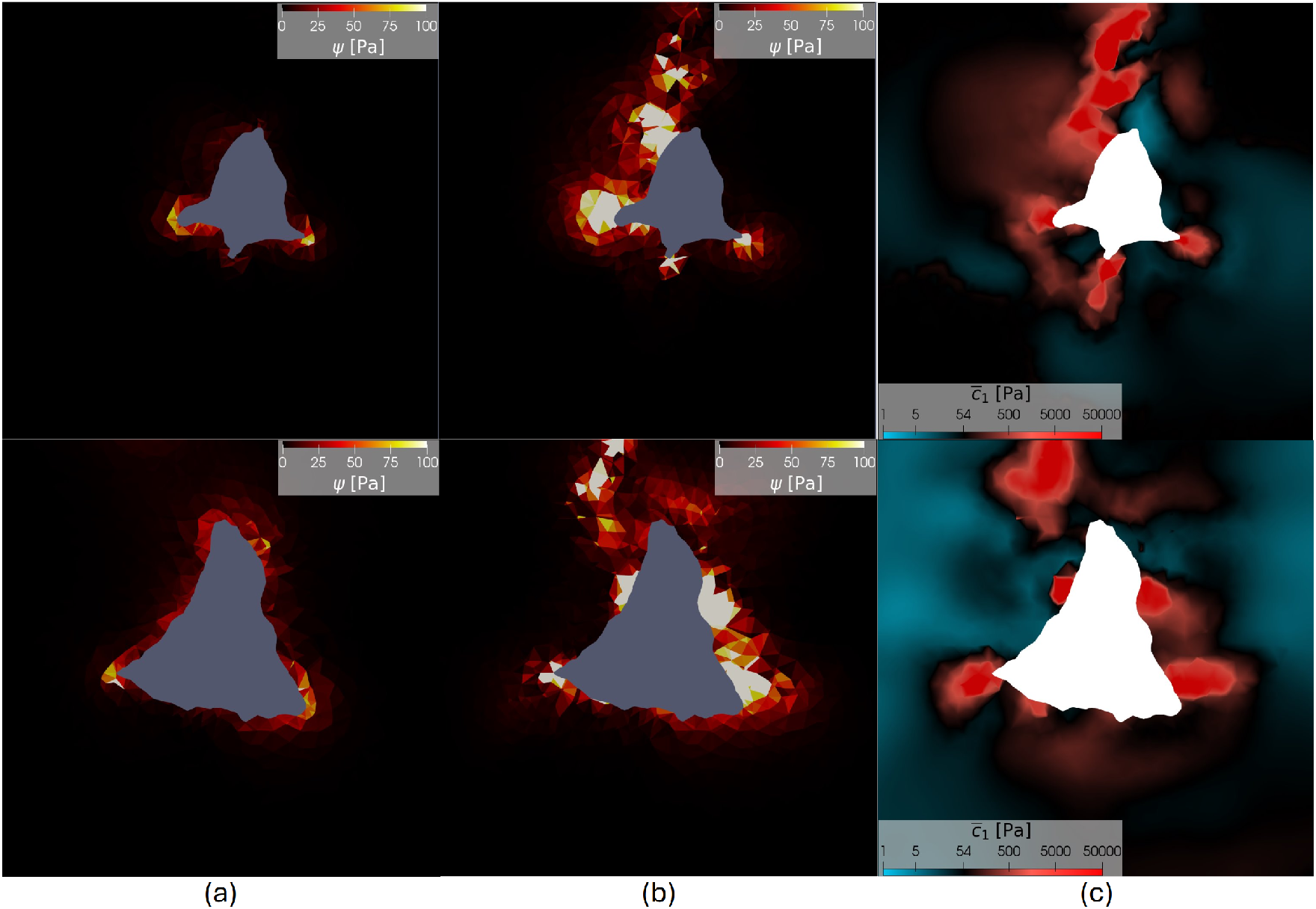
Comparisons of strain-energy density Ψ assuming homogeneous vs. heterogeneous modulus demonstrated large differences in simulated cellular mechanical behavior. MVICs are presented per-row in the same order as in Figure 7. We render element-wise projected Ψ and predicted nodal 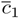 on representative cross-sections. The form of strain-energy density is given in Equation 1. The scale has been truncated to highlight a representative range of variation. In (a)-(b), the hollow inclusion of the cell is in gray. (a) Ψ assuming hydrogel modulus is unmodified, *i*.*e*. 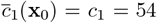 Pa the far-field value uniformly. (b) Ψ computed with inverse-model-output 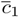. (c) Inverse-model-output 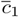. The hollow inclusion of the cell is in white, unmodified hydrogel modulus (54 Pa) is in black, degraded hydrogel modulus is in blue, and stiffened hydrogel modulus is in red.

Simulated displacements and *J* values predicted by the inverse model matched the target experimental quantities very well (Figure 10). The simulated standard deviations of *J* were 0.0385 and 0.0295 respectively, close to target standard deviations 0.0432 and 0.0364. The differences between these standard deviations were below the level of the noise in the data, which we estimated contributed 0.009 error based on the realistic test problem (Section 3.1.2). This match between experimental *J* and simulated *J* demonstrated consistent modeling of compressibility.

**Figure 10.**
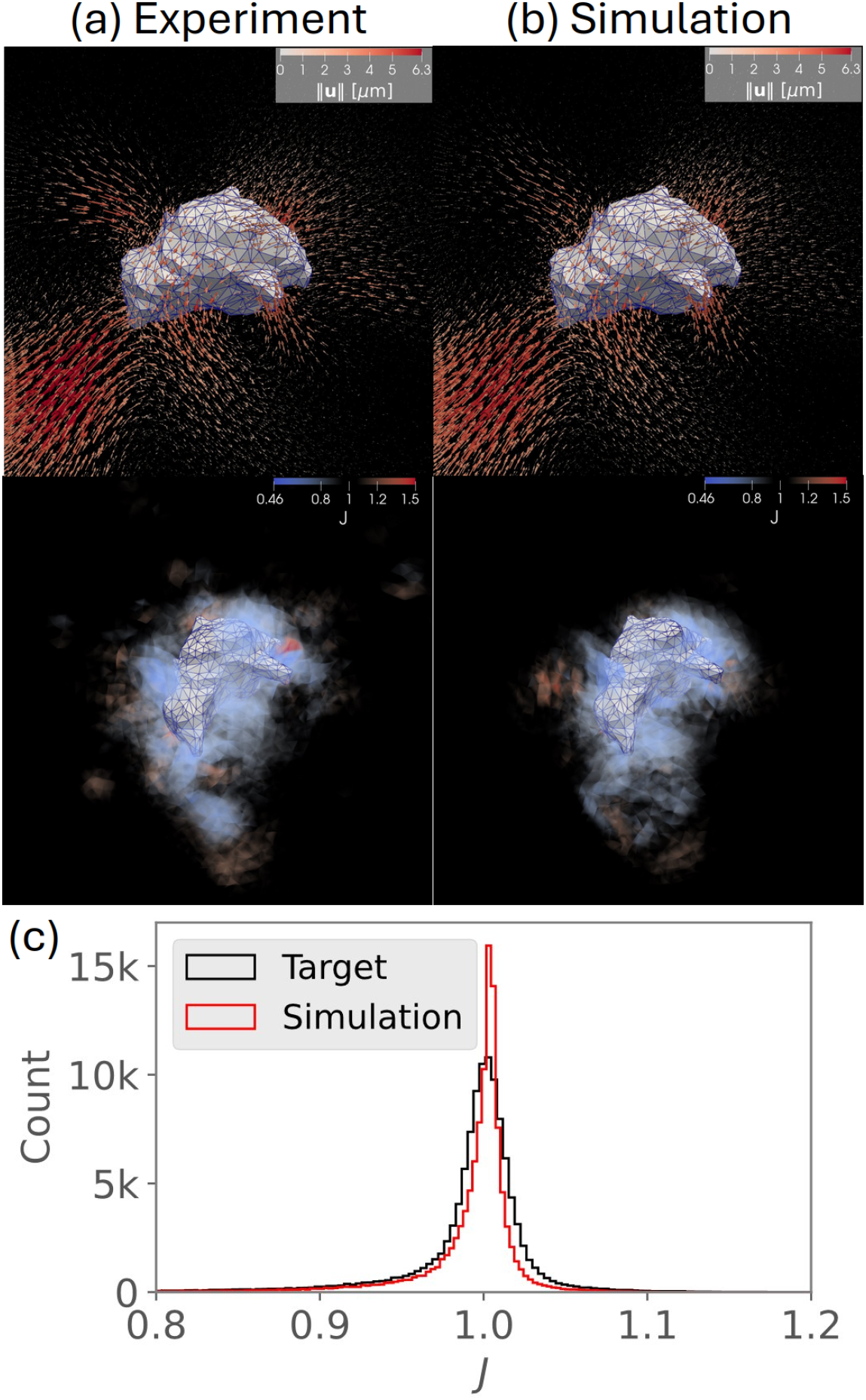
Target experimental displacements *J* vs. inverse-model prediction for a representative cell. Gray mesh is the cellular surface, shaded arrows in the top row are displacements and shaded transparent elements in the middle row are *J*. The top row is in the current configuration while the middle row is in the referential configuration. (a) Target kinematic quantities from GPR-interpolation. (b) Inverse-model-simulated kinematic quantities from the forward model with the predicted modulus field. (c) Comparison of element-based *J* values within the event horizon between target and inverse-model prediction.

## 5 DISCUSSION

### 5.1 Overview of Major Findings

In the present work we developed a novel inverse-modeling pipeline capable of handling compressible hydrogel behaviors for large-deformation 3D TFM analysis. We utilized a number of modifications in our approach to ensure numerical stability as well as to accurately account for the spatial heterogeneity in effective hydrogel moduli. We extensively validated our modeling pipeline *via* synthetic test cases featuring realistic cell geometries, displacement fields, and noise conditions. Importantly, we found that accurately accounting for hydrogel compressibility was critical for accurate capture of the spatial variation in modulus as well as traction-force determinations. Our test cases clearly demonstrated the necessity of numerical stabilization. When the regularization parameter started too small, numerical instability caused the inverse model to diverge. Our previous approach only provided for using a single regularization parameter, contributing to its limitation in recovering only modest modulus modifications. Similarly, the improved choice of *L*^2^ Riesz representation in gradient computations was required for convergence of the initial stages. Whereas alternative inverse-modeling approaches to TFM make simplifications such as treating the system as 2D, using linear elasticity, and assuming homogeneity [5–8, 11–13], our approach achieved successful recovery in the full-3D nonlinear heterogeneous material-modeling context.

Our inverse-modeling approach achieved high fidelity modulus and compressibility recovery even in the presence of noisy displacements, handled using Tikhonov regularization. After establishing validity under realistic conditions and matching levels of compressibility between ground truth and inverse model, we then established that proper modeling of compressibility was a key component that to our knowledge had not previously been rigorously evaluated for the case of large deformations in 3D TFM. Previous studies considering the influence of compressibility of hydrogels in TFM have been restricted to homogeneous linear-elastic models for small deformation, in which case it was demonstrated that the Poisson ratio must be properly calibrated [44]. These previous findings for infinitesimal-deformation models are not relevant to the case of actual large deformations in hydrogels. In our synthetic test cases where a less-compressible material model was exercised, the modulus field was poorly recovered. In particular, improperly accounting for compressibility biased the model to predict stiffening not present in the target. Likewise, predicted strain energy and traction forces contained large errors.

We next demonstrated the utility of our approach by analyzing novel high-fidelity TFM experimental data using human MVICs embedded in PEG hydrogels, the latter of which are commonly studied in TFM. Values of *J* = 1*±*0.6 in the vicinity of the cell were observed, establishing the presence of compressible behavior in PEG hydrogels. It is also known that MVICs exhibit high metabolic activities [45–47], and thus are likely to induce very large modifications to hydrogel modulus. Indeed, our approach detected local modifications of hydrogel moduli spanning 3.6 Pa to 2.4 MPa, much wider than than the range detectable by our previous approach, 5.4 Pa to 540 Pa [14]. Moreover, we found that accounting for the full range of modulus yielded predictions of cellular mechanical behavior different by more than an order of magnitude from the homogeneity assumption. Thereby the example of human MVICs demonstrated the applicability of our approach accounting for both compressibility and extensive heterogeneity in cellularized hydrogels.

The exact mechanisms underlying the observed compressiblity remain unknown, but are most likely due to the intrinsic poroelasticity of hydrogels [48]. Due to imaging time limitations, it is not possible to obtain dynamic information in the TFM experiment. Specially, in the time period between collecting image stacks, the system settled into a steady-state, and poroelastic fluid flows occured, resulting in local compression and expansion. In this case, those changes would correspond to fluid exudation and swelling respectively. Our modeling the hydrogel as elastic remains valid whether the hydrogel is in fact poroelastic or viscoelastic, as any time-dependent effects have been dissipated.

### 5.2 Limitations and Future Directions

The experimental 3D TFM approach that our method is designed for is limited to imaging the cell and hydrogel at only two time points. Chemical treatment, including cytochalasin-D, takes a long time period, 30 minutes, to progressively alter cell stress-fiber force generation. Collecting *z*-stacks of microscope images also takes time, and long exposure to lasers can cause photo-bleaching of cells. Consequently, we consider only a single displacement field in our approach. Cellular remodeling of the hydrogel may also be partly responsible for altering its compressibility, which our method does not take into account; we only model *c*_1_(**x**_0_) as spatially-varying while *D*_1_ is homogeneous. Our method also cannot detect possible anisotropy of ECM-infused hydrogel, which would possibly be associated with orientation of fibers such as collagen and elastin [49]. Current studies are underway to address these potential effects. We have previously developed a method to estimate internal cell stress fiber force levels and orientations [50]. This process required inverse modeling to determine heterogeneous hydrogel properties as a prerequisite. Adapting the stress-fiber inverse model to our new material model formulation in future work will allow more accurate insights into internal cell organization and activity. Simultaneous determination of stress-fiber and ECM structure will then allow detailed comparisons of diseased and healthy phenotypes of cells. This work may also be extended to employ other hyperelastic material models by adapting them with a similar exponential modulus representation.

## 7. AUTHOR CONTRIBUTIONS – CRediT

**Gabriel Peery:** Methodology (lead), Software (lead), Validation (lead), Formal Analysis (lead), Investigation (lead), Data Curation (equal), Writing – Original Draft (lead), Visualization (lead), Funding Acquisition (equal). **Toni M. West:** Software (supporting), Investigation (supporting), Data Curation (equal), Writing – Review & Editing (supporting), Funding Acquisition (equal). **Sanjana S. Chemuturi:** Data Curation (equal), Formal Analysis (supporting), Writing – Review & Editing (supporting). **Jodie H. Pham:** Data Curation (equal), Formal Analysis (supporting), Writing – Review & Editing (supporting). **Giovanni Ferrari:** Resources (equal), Funding Acquisition (equal), Data Curation (equal), Writing – Review & Editing (supporting). **Michael S. Sacks:** Conceptualization (lead), Methodology (supporting), Resources (equal), Writing – Review & Editing (lead), Supervision (lead), Project Administration (lead), Funding Acquisition (equal).

## 8. ACKNOWLEDGMENTS

We would like to thank Giovanni Ferrari’s group for providing the MVICs for this study.

## 10. CODE AND DATA AVAILABILITY

Our implementation of this inverse modeling approach is available on GitHub:

https://github.com/WCCMS-UTAustin/3DTFM.

Data involved in the synthetic test cases is publicly available at Mendeley Data [51]: DOI: 10.17632/nmfxvtbrbd.1

Data involved in the Human MVIC examples is available upon reasonable request.

## 11. ETHICS STATEMENT

Human MVICs were isolated from consented patients undergoing mitral valve replacement by the Columbia Biobank for Translational Science (IRB-AAAR6796). The de-identified MVICs were then shipped to the University of Texas where culturing and imaging occurred (IBC-2023-00293).

